# Thinking like a naturalist: enhancing computer vision of citizen science images by harnessing contextual data

**DOI:** 10.1101/730887

**Authors:** J. Christopher D. Terry, Helen E. Roy, Tom A. August

## Abstract

1. The accurate identification of species in images submitted by citizen scientists is currently a bottleneck for many data uses. Machine learning tools offer the potential to provide rapid, objective and scalable species identification for the benefit of many aspects of ecological science. Currently, most approaches only make use of image pixel data for classification. However, an experienced naturalist would also use a wide variety of contextual information such as the location and date of recording.
2. Here, we examine the automated identification of ladybird (Coccinellidae) records from the British Isles submitted to the UK Ladybird Survey, a volunteer-led mass participation recording scheme. Each image is associated with metadata; a date, location and recorder ID, which can be cross-referenced with other data sources to determine local weather at the time of recording, habitat types and the experience of the observer. We built multi-input neural network models that synthesise metadata and images to identify records to species level.
3. We show that machine learning models can effectively harness contextual information to improve the interpretation of images. Against an image-only baseline of 48.2%, we observe a 9.1 percentage-point improvement in top-1 accuracy with a multi-input model compared to only a 3.6% increase when using an ensemble of image and metadata models. This suggests that contextual data is being used to interpret an image, beyond just providing a prior expectation. We show that our neural network models appear to be utilising similar pieces of evidence as human naturalists to make identifications.
4. Metadata is a key tool for human naturalists. We show it can also be harnessed by computer vision systems. Contextualisation offers considerable extra information, particularly for challenging species, even within small and relatively homogeneous areas such as the British Isles. Although complex relationships between disparate sources of information can be profitably interpreted by simple neural network architectures, there is likely considerable room for further progress. Contextualising images has the potential to lead to a step change in the accuracy of automated identification tools, with considerable benefits for large scale verification of submitted records.

## Introduction

Large-scale and accurate biodiversity monitoring is a cornerstone of understanding ecosystems and human impacts upon them (IPBES, 2019). Recent advances in artificial intelligence have revolutionised the outlook for automated tools to provide rapid, scalable, objective and accurate species identification and enumeration (Wäldchen & Mäder, 2018; Weinstein, 2018; Torney et al., 2019; Willi et al., 2019). Improved accuracy levels could revolutionise the capacity of biodiversity monitoring and invasive species surveillance programs (August et al., 2015). Nonetheless, at present, general-purpose automated classification of animal species is currently some distance from the level of accuracy obtained by humans, and the potential remains underutilised.

The large data requirements and capacity of machine learning has led to a close association with citizen science projects (Wäldchen & Mäder, 2018), where volunteers contribute scientific data (Silvertown, 2009). Citizen scientists can accurately crowd-source identification of researcher-gathered images (e.g. Snapshot Serengeti; Swanson et al., 2015), generate records to be validated by experts (e.g. iRecord; Pocock, Roy, Preston, & Roy, 2015) or both simultaneously (e.g. iNaturalist; iNaturalist.org). However, there can be a considerable lag between record submission and human verification. If computer vision tools could generate more rapid, or even instantaneous, identifications it could assist with citizen scientist recruitment and retention. While image acquisition by researchers can be directly controlled and lead to high accuracies (Rzanny, Seeland, Wäldchen, & Mäder, 2017; Marques et al., 2018), images from citizen science projects are highly variable and pose considerable challenges for computer vision (Van Horn et al., 2017).

Most automatic species identification tools only make use of images (Weinstein 2018). However, an experienced naturalist would utilise a wide variety of contextual information when making an identification. This is particularly the case when distinguishing ‘difficult’ species, where background information about the record may be essential for a confident identification. In a machine learning context, this supplementary information about an image (metadata) can be split into two categories (Figure 1). Primary metadata is directly associated with a record such as GPS-coordinates, date of recording and the identity of the recorder. Derived (secondary) metadata is generated through cross-referencing with other sources of information to place this metadata into a more informative context (Tang, Paluri, Fei-Fei, Fergus, & Bourdev, 2015). In an ecological context, this may include weather records, maps of species distribution, climate or habitat, phenology records, recorder experience, or any other information source that could support an identification.

**Figure 1.**
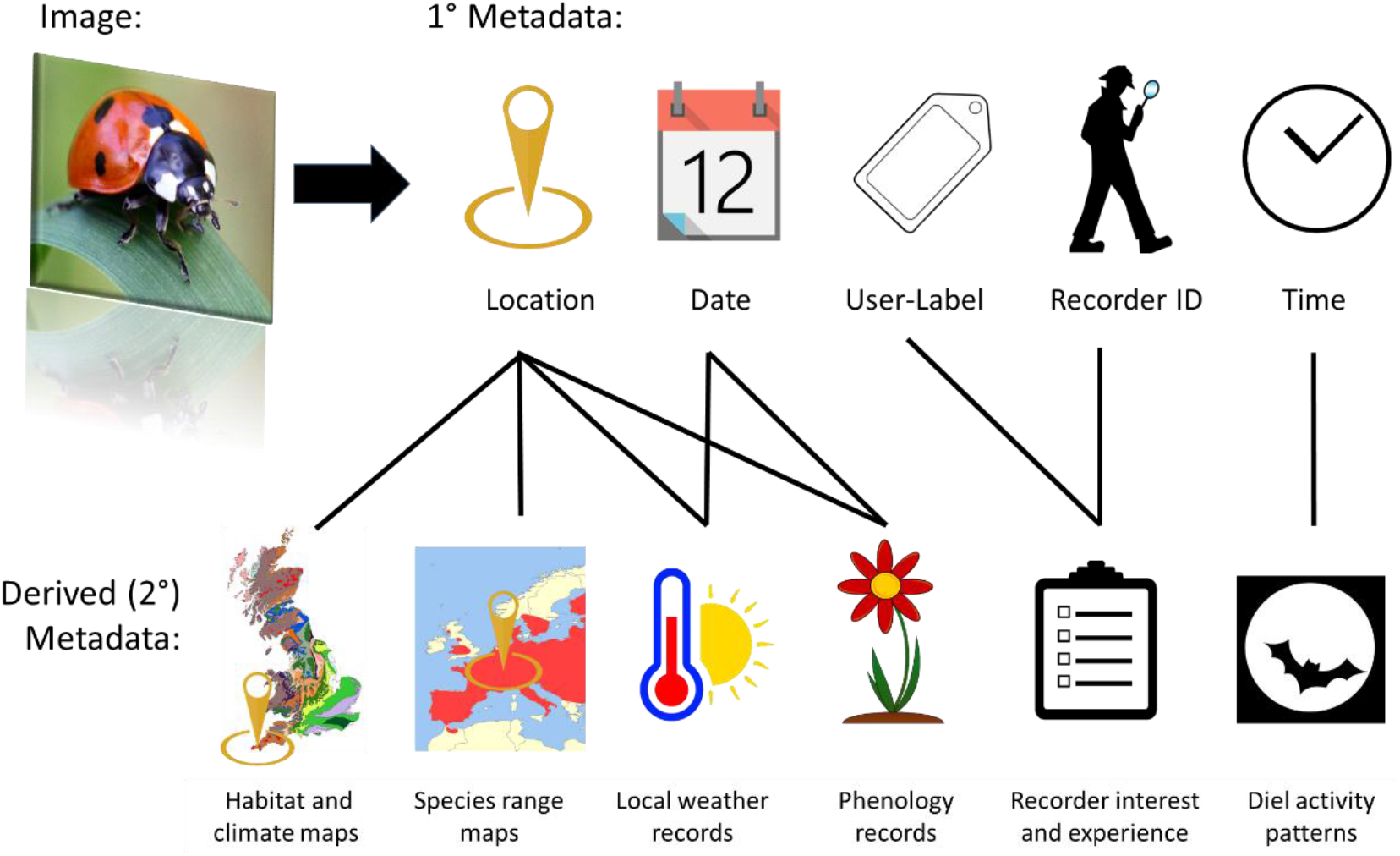
Relationships between categories of metadata. Primary metadata are basic attributes of the record directly associated with an image such as the date or location. By contrast, derived (or secondary) metadata requires cross-reference to external databases, which may include physical, ecological or social data. External sources of information may be fixed and stable (such as habitat maps) or dynamic and require updating in order to keep the model up to date (such as weather records or recorder experience).

Efforts to include contextual spatio-temporal information have largely focused on reducing the list of potential species that may be expected in a given area. iRecord (www.brc.ac.uk/irecord) partially automates this process, flagging records to expert verifiers that are labelled as being outside of the known range. Distribution priors have been shown to be effective in improving the identification of North American birds (Berg et al., 2014), images in the iNaturalist dataset (Mac Aodha, Cole, & Perona, 2019) and generating location-specific shortlists of German plants (Wittich, Seeland, Wäldchen, Rzanny, & Mäder, 2018). This approach can greatly reduce the risk of non-sensical identifications that otherwise lead to considerable scepticism over the use of automated methods (Gaston & O’Neill, 2004). Nevertheless, this ‘filtering’ approach does not make full use the available data. Many species vary in appearance seasonally or across their range. For example, the proportion of the melanic form of the 2-spot ladybird *Adalia bipunctata* varies greatly across the UK (Creed, 1966). To an expert naturalist, metadata can do more than shorten the list of potential identifications - it can help to interpret the image itself. For example, juveniles, flowers or breeding plumage may only be observed in narrow time windows or there may be geographic variation in colour patterns. Consequently, certain features within an image (e.g. spots on a butterfly’s wing) may only aid in determining a species in specific regions, or times of year. It would only be worth looking for a particular pattern when that species and lifestage is active. Synthesising and making use of such disparate sets of information is challenging for humans even when detailed data is available, and such expertise requires many years to build. By contrast, neural networks are ideally suited to drawing together diverse sources in such a way to gain the maximal amount of information.

Ladybirds (Coleoptera: Coccinellidae) are a charismatic insect family that garner substantial public interest, with large numbers of submitted records to citizen science monitoring schemes around the world (Gardiner et al., 2012). Identification of ladybirds is challenging for both human (Jouveau, Delaunay, Vignes-Lebbe, & Nattier, 2018) and artificial intelligence (Van Horn et al., 2017) because of a number of morphological features. Many species of ladybird have polymorphic elytral colour patterns, with some species seemingly mimicking others, and so are principally disambiguated by size. However, size is extremely challenging for artificial intelligence to automatically infer from a single image without standardised scales (Laina, Rupprecht, Belagiannis, Tombari, & Navab, 2016). As an example the invasive Harlequin ladybird *Harmonia axyridis* (which has been a particular focus for research, Roy et al., 2016), is a polymorphic species and can resemble a number of other species. Consequently, the Harlequin ladybird is frequently misidentified by citizen scientists (Gardiner et al., 2012) but can be distinguished on the basis of its large size. Currently, submissions to the UK Ladybird Survey (www.ladybird-survey.org) are managed by a small number of expert verifiers, imposing a large burden on the expert community. There is growing interest in expanding the geographic scope of the survey with the recent launch of a smartphone app for recording ladybirds across Europe (https://european-ladybirds.brc.ac.uk/). The UK ladybird survey (and associated European extension) therefore represents an example of a programme where a reliable automated identification tool could help to increase the use of citizen science to document biodiversity across the globe.

Classification tools that only use image data are not making maximal use of the information available to human experts. Here we demonstrate methods to incorporate metadata directly within neural networks used for the classification of images of ladybirds submitted to the UK Ladybird Survey. We examine if metadata can significantly improve classification accuracy, thereby increasing their potential to assist in large-scale biodiversity monitoring, by:

1. Comparing the classification accuracy of classifiers incorporating metadata compared to image-only classifiers.
2. Exploring whether neural networks make use of the same pieces of metadata information that a human experts do.

## Methods

### Data

Records of ladybirds (Coccinellidae) were sourced from the UK Biological Records Centre (www.brc.ac.uk). These were filtered to include only those from within the British Isles, from 2013 to 2018 inclusive, that contained an image and had been verified by an expert assessor. Records were distributed across the whole of the British Isles, although records were more frequent near more-heavily populated areas (Figure S1). The date range was selected based on a notable increase in records from 2013 with the release of a mobile app (iRecord Ladybirds). Identifications of records by expert verifiers was based on uploaded images and associated information including the species determination of the original observer, location, date, associated comments and (where known) the degree of skill of the recorder.

Of the 47 species of ladybird that had been recorded at least once in the UK (Duff, 2018), only 18 species (listed in table 1) had at least 170 usable records, which we took as our lower cut-off to ensure each species was represented by at least 120 unique training images. We judged that fewer training images would not result in accurate classification. These 18 species made up 97% of the total ladybird records during 2013-2018. Even after removing species with fewer than 170 usable records, the data set is highly imbalanced (Table 1), with two species making up the bulk of records: 7-spot ladybird *Coccinella septempunctata* (25.8%) and the highly polymorphic Harlequin ladybird (44.5%).

### Images

Records were manually scanned to remove the majority of images predominantly of eggs, larvae or pupae, ‘contextual’ images of habitat area, images including multiple species, and images that had been uploaded repeatedly. Larval and pupal images were overwhelming dominated by the highly distinctive Harlequin ladybird larvae or pupae (78%). Where a single record had multiple associated images, only the first was used. Images were centre cropped to square and then rescaled to 299×299 pixels. Example images for each species are shown in Figure 2. After all data cleaning steps, the dataset had 39,877 records in total.

**Figure 2.**
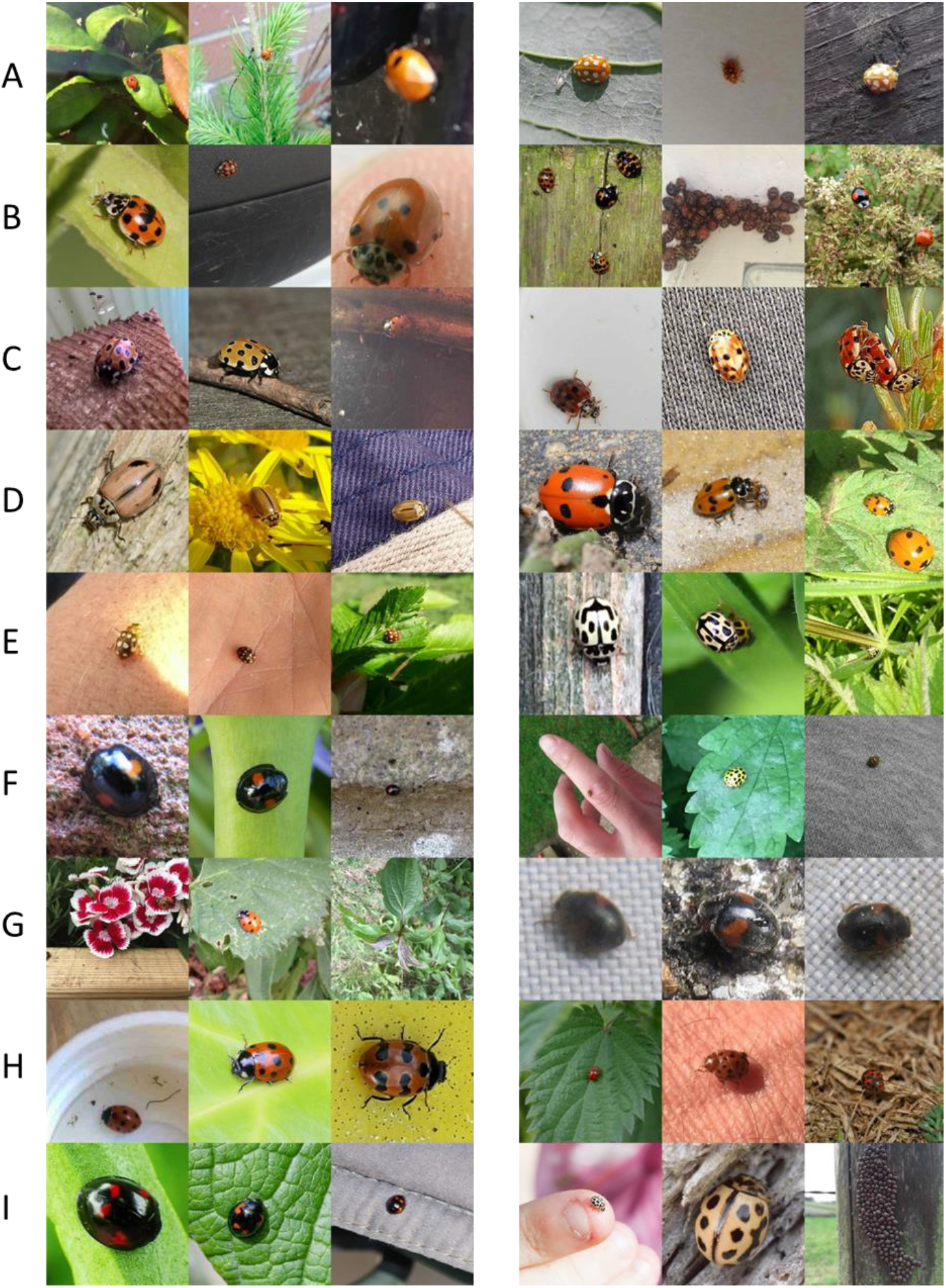
Three randomly selected images from each of the 18 ladybird species in our dataset, demonstrating the wide variety of poses, sizes and backgrounds. Images have been centre cropped to square and resized to 299×299. Species are listed alphabetically: Left column: a) *Adalia bipunctata*, b) *Adalia decempunctata*, c) *Anatis ocellata*, d) *Aphidecta obliterata*, e) *Calvia quattuordecimguttata*, f) *Chilocorus renipustulatus*, g) *Coccinella septempunctata*, h) *Coccinella undecimpunctata*, i) *Exochomus quadripustulatus*. Right column: a) *Halyzia sedecimguttata*, b) *Harmonia axyridis*, c) *Harmonia quadripunctata*, d) *Hippodamia variegata*, e) *Propylea quattrodecimpunctata*, f) *Psyllobora vigintiduopunctata*, g) *Scymnus interruptus*, h) *Subcoccinella vigintiquattropunctata*, i) *Tytthaspis sedecimpunctata*.

### Metadata

We constructed models that made use of different subsets of the available metadata. The first (the primary metadata model) took only three pieces of primary metadata, drawn directly from the UK Ladybird Survey dataset: longitude, latitude and date. We represented date by day-of-year, excluding year values since information on ‘year’ would not be transferable to future records. The second model (the derived metadata model) supplemented the primary metadata with secondary metadata: data generated with additional reference to external sources of information, namely weather records, habitat and recorder expertise. We did not use the original citizen scientist species determination in our models, since it was too powerful compared to other sources of information (correct over 92% of the time) and did not align with the goal of fully automated identification.

Temperature records were accessed from the Midas database (Met Office, 2012), selecting data from the 88 UK stations with fewer than 20 missing records (2013 to 2018). Occasional missing values were imputed with a polynomial spline. Using the closest weather station to the record, maximum daily temperature for each day in the 14 preceding days (*d-1:d-15*) and weekly average maximum daily temperatures for each of the 8 weeks preceding the high resolution period (*d-16:d-71*) were accessed.

Local habitat information was derived from a 1km resolution land cover map (Rowland et al., 2017). This provides percentages in each 1km grid of 21 target habitat classes (e.g. ‘urban’, ‘coniferous woodland’, ‘heather’, etc.). Where no data was available, each habitat was assumed to be 0.

We calculated a ‘recorder experience’ variable as the cumulative count of records submitted by that recorder at the time of each record. Only records of ladybirds in our dataset were included in this count. Where no unique recorder ID was available, that record was assumed to be a first record.

This led to a one-dimensional metadata vector of length 47 (day-of-year, latitude, longitude, 14 daily maximum temperature records, 8 weekly average temperature records, 21 habitat frequencies and recorder experience) associated with each image.

### Machine learning model architecture

We built and fit convolutional neural network models (Goodfellow, Bengio, & Courville, 2016) in R 3.5.3 using the functional model framework of the *keras* package (Allaire & Chollet, 2019). We used the TensorFlow backend on a Nvidia GTX 1080 Ti GPU. R code used to train the models is available at *github.com/jcdterry/LadybirdID_Public* and the core model architecture code is summarised in SI. We first constructed and trained image-only and metadata-only models. Once these had separately attained maximum performance, these were then combined to form the core of a multi-input model that takes both an image and metadata as input variables. For all models we conducted extensive hyperparameter searches to determine model architecture, extent of data-augmentation, regularisation parameters, learning rates and training times.

A schematic of the model architectures is shown in Figure 3. The metadata models were built with a simple architecture of two densely connected layers and a softmax classifier layer. For the image-model, the Inception-ResNet-v2 architecture (Szegedy, Ioffe, Vanhoucke, & Alemi, 2016) was used as an initial feature extractor. This is a very deep architecture that had been pretrained on the large imageNet dataset to extract meaningful features from a generic set of images. This transfer learning approach greatly expedites the training process and has previously achieved high accuracy in tests on the iNaturalist data set of citizen science records (e.g. Cui, Song, Sun, Howard, & Belongie, 2018) and for the identification of insects (Martineau et al 2018). To repurpose the model, we replaced the imageNet classification layer with new layers and trained the model on our dataset. The combined model was built by removing the classifier layers from the metadata and image models, concatenating the two outputs, and adding further layers. This fusion approach has been successfully used in the categorisation of satellite data (Minetto & Segundo 2019).

**Figure 3.**
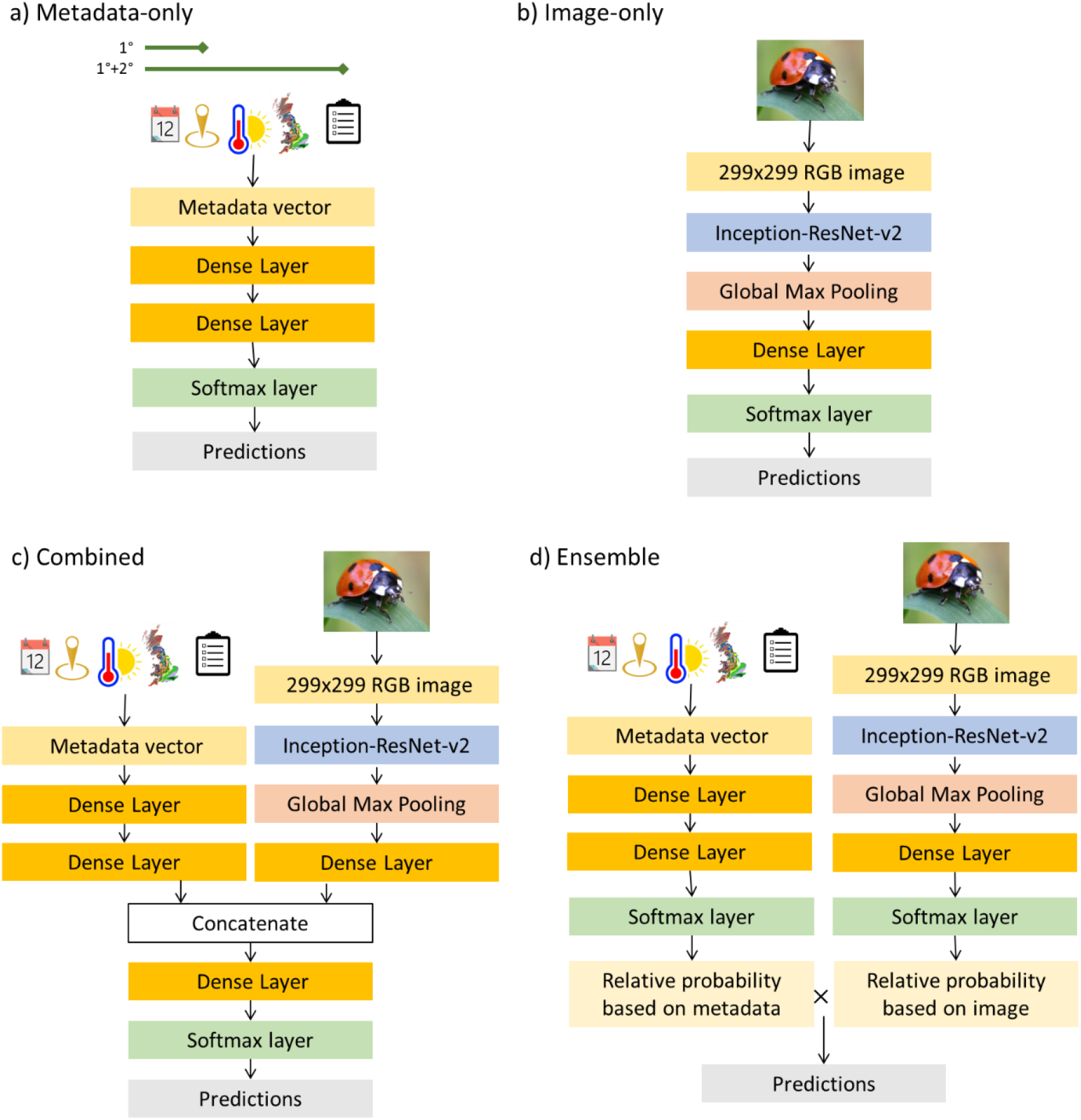
Outline schematic of the difference in model architectures. Dense layers are the principle component of neural networks, that fit linkages between every input and output node. All our dense layers incorporated a rectified linear unit (ReLU) non-linear activation function. Inception-ResNet-v2 is a very deep feature extraction model incorporating many convolutional layers and originally trained to classify a diverse set of objects, that we refined by retraining on our ladybird dataset. The global max pooling stage summarises the outputs of the image feature extractor for further computation by dense layers. Softmax layers output a vector that sums to one, which can be interpreted as probabilities of each potential category. Dropout, noise, batch normalisation and other regularisation features enacted only during training time are not shown here for simplicity. R code to build models using the *keras* R package (Allaire & Chollet, 2019) is given in SI, which also details further hyperparameters such as the size of the each layer.

### Model Training

Species records in the UK Ladybird Survey, like most biological record datasets (Van Horn et al., 2017), are highly skewed towards certain common species (Table 1). As predictive models are not perfect, such class-imbalanced data leads to critical choices about how to best assess ‘accuracy’. Overall accuracy may be maximised by rarely or never assigning species to unusual categories. A citizen scientist may prefer the maximum accuracy for the species in front of them (which is likely to be a commonly reported species). However, in an ecological science context, rare (or more precisely, rarely reported) species are often of particular interest to researchers managing citizen science projects.

The total dataset was randomly partitioned into training (70%), validation (15%) and test (15%) sets. To address the class-imbalance, we followed the approach suggested by Buda, Maki, & Mazurowski, (2018) and re-balanced our training set through up-sampling and down-sampling the available records. We did this so that each species had 2000 effective training records. Consequently, our underlying models did not have direct access to the information that, all else being equal, certain species are far more likely than others. This reduces the potential for the model ‘cheating’ during training by fixating on common species and ignoring rare species. To demonstrate the potential to improve overall accuracy by taking into account the relative frequency of each species, we tested weighted versions of each of the models. In these, the relative probability assigned to each species from each unweighted model (*P_i_*) were scaled by the relative frequency of each of the species (*F_i_*) in the training data as: *P_weighted_i__*. ∝ *P_i_ F_i_*.

To reduce overfitting, we made extensive use of image augmentation, weight regularisation, batch normalisation, dropout layers during training and introduced Gaussian noise on the metadata vector. Training optimisation was based on a categorical cross-entropy loss function using the *‘Adam’* adaptive moment estimation optimiser (Kingma & Ba, 2014). During training, if validation loss had reached a plateau, learning rate was reduced automatically. Training was stopped (and the best model restored) if there had been no further improvement in validation loss over at least four epochs.

After fitting the derived metadata, image-only and combined models, a simple ensemble model taking a weighted average of the derived metadata and image-only model predictions was also constructed and tested. This could be considered equivalent to using the metadata to construct a prior expectation for the predictions of the image model:

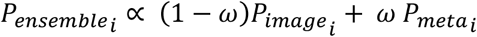

where the weighting (*ω*) between the metadata and image model probabilities was determined by optimising the ensemble model top-1 accuracy on the validation set.

### Model Testing and Evaluation

Overall and species-level model performance was assessed in terms of top-1 (was the true ID rated most likely) and top-3 (was the true ID amongst the three options rated most highly) accuracy. Because model accuracy will be dependent on the split of data into testing and training sets, and because model optimisation is a non-deterministic process, we repeated the entire model fitting process 5 times. For each repeat, assignment of images to training, validation and test sets was randomised.

### Role of Metadata Components

To examine the dependence of the model on each aspect of the metadata we examined the decline in top-3 accuracy for each species when elements of metadata were randomised by reshuffling sets of values within the test set. We did this separately for the spatial coordinates, day–of-year, temperatures data, habitats data and recorder expertise.

## Results

Across each of our training-test split realisations, combined multi-input models showed a marked and consistent improvement on both the image-only (+ 9.1 percentage points) and the ensemble models (+ 3.6 percentage points) (Figure 4). Species-level accuracies (averaged across the 5 split realisations) for each of the models are reported in Table 1. There was no correlation between the species-specific accuracy of the metadata-only model and the image-only model (Spearman’s rank correlation test *ρ* =0.23, p=0.34). There was, however, a strong correlation at a per-species level between the fraction correctly identified by the original citizen-scientist recorder and the combined model (*ρ* = 0.65, p = 0.003).

**Figure 4.**
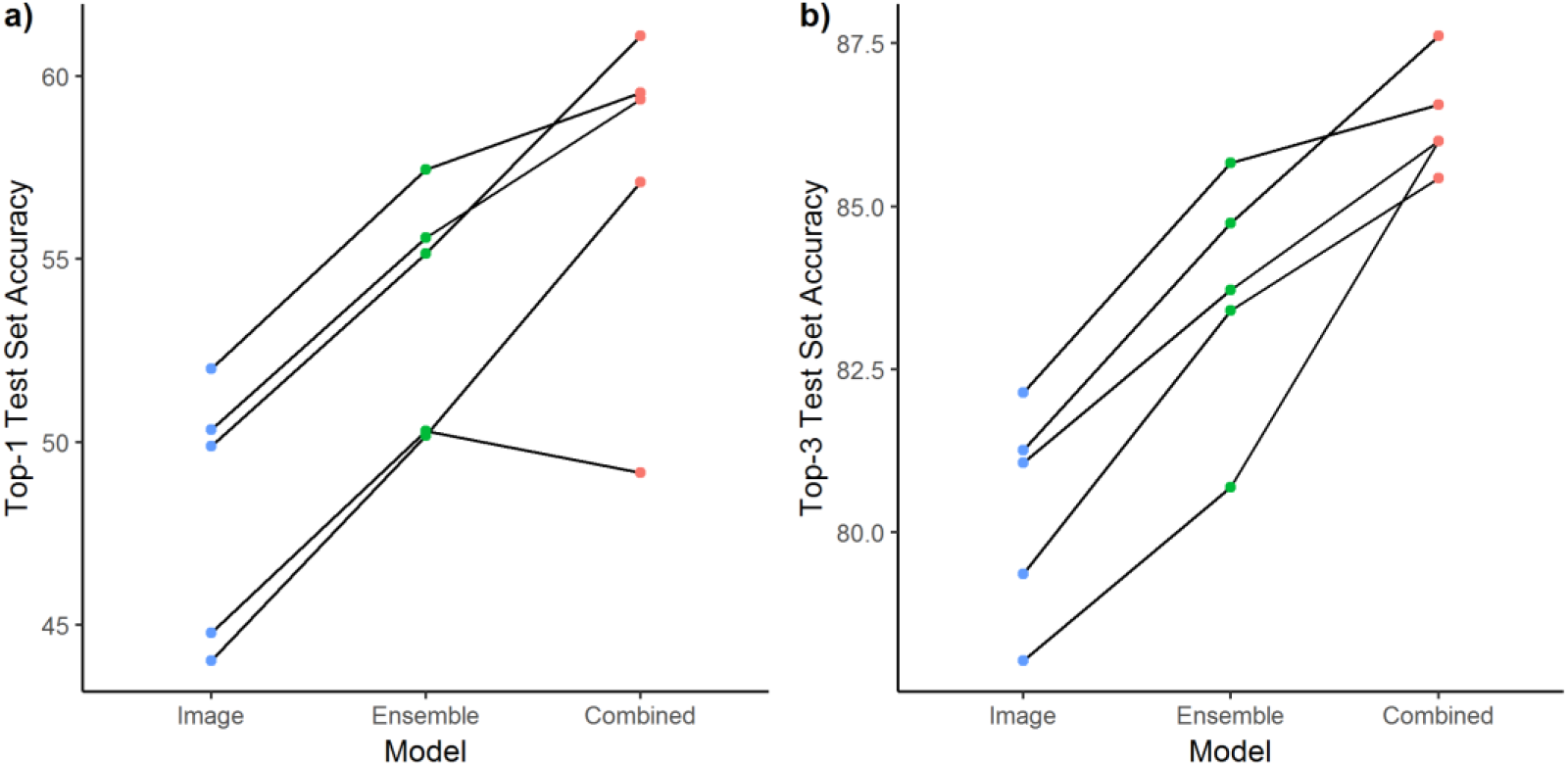
Consistent improvement in top-1 (a) and top-3 (b) accuracy from image only models to models with the incorporation of metadata. An image-only model can be improved by ensembling with a metadata model, but further improvements can be gained from fitting combined multi-input models. Lines show 5 suites of models trained on a different train-validation-test randomisations.

**Table 1.** Average per-species top-1 accuracy across the suite of models. Citizen scientist accuracy is determined by frequency by which the label assigned by the recorder corresponds to the verified species name. Equivalent tables for top-3 accuracy and for accuracy including a prior weighting based on relative frequency are given in SI.

| Species | Relative Frequency | Citizen Scientist | Metadata Only |  | Image Only | Image and Metadata |  |
| --- | --- | --- | --- | --- | --- | --- | --- |
|  |  |  | Primary | Derived |  | Combined | Ensemble |
| **Overall** |  | 92.4 | 15.9 | 22.4 | 48.2 | 57.3 | 53.7 |
| *Adalia bipunctata* | 5.3 | 97.3 | 10.9 | 22.5 | 56.4 | 58.9 | 58.5 |
| *Adalia decempunctata* | 2.9 | 85.8 | 1.9 | 2.2 | 24.6 | 22.9 | 23.3 |
| *Anatis ocellata* | 0.5 | 94.5 | 2.7 | 15.3 | 37.3 | 41.3 | 42.0 |
| *Aphidecta oblitterata* | 0.7 | 96.5 | 63.2 | 55.3 | 71.6 | 80.0 | 81.6 |
| *Calvia quattuordecimguttata* | 1.8 | 92.5 | 0.2 | 3.4 | 70.0 | 55.8 | 69.8 |
| *Chilocorus renipustulatus* | 1.2 | 93.2 | 3.1 | 16.9 | 47.6 | 47.0 | 49.6 |
| *Coccinella septempunctata* | 26.1 | 95.5 | 0.0 | 0.3 | 64.8 | 64.2 | 62.9 |
| *Coccinella undecimpunctata* | 0.7 | 94.0 | 5.1 | 27.2 | 58.5 | 58.5 | 62.1 |
| *Exochomus quadripustulatus* | 1.5 | 92.0 | 15.2 | 26.9 | 37.9 | 43.9 | 40.0 |
| *Halyzia sedecimguttata* | 3.7 | 93.9 | 0.8 | 7.7 | 65.6 | 73.5 | 66.1 |
| *Harmonia axyridis* | 44.1 | 89.6 | 27.9 | 38.6 | 34.7 | 53.6 | 47.1 |
| *Harmonia quadripunctata* | 0.4 | 94.3 | 6.2 | 12.3 | 37.7 | 43.1 | 39.2 |
| *Hippodamia variegata* | 0.6 | 93.2 | 35.4 | 32.0 | 28.0 | 46.9 | 38.9 |
| *Propylea quatuordecimpunctata* | 4.5 | 94.8 | 28.1 | 22.3 | 58.6 | 62.7 | 59.3 |
| *Psyllobora vigintiduopunctata* | 3.0 | 98.5 | 5.2 | 11.8 | 56.3 | 58.5 | 58.1 |
| *Scymnus interruptus* | 0.4 | 98.2 | 93.6 | 76.0 | 88.0 | 89.6 | 90.4 |
| *Subcoccinella vigintiquatuor punctata* | 1.6 | 96.2 | 6.2 | 17.8 | 62.6 | 67.3 | 64.5 |
| *Tytthaspis sedecimpunctata* | 1.0 | 91.2 | 7.0 | 21.9 | 43.5 | 51.1 | 50.8 |

The overall accuracy of all models could be greatly improved by weighting the output probabilities by the prior expectation given the relative frequency of each species. For example, the average top-1 accuracy of the combined model rises from 57% to 69%. However, these gains are made at the cost of very infrequently identifying unusual species correctly. With a weighted model the two most commonly observed species, Harlequin and Seven-spot ladybirds, are correctly identified 90% and 89% of the time respectively. However, 12 infrequently observed species are correctly identified in less than 12% of cases.

The derived metadata model had an overall top-3 accuracy of 43.7% and was making at least some use of all the components of the metadata since randomising each group caused a decline in accuracy. Accuracy of the metadata-only model peaked spatially away from the south-east of the British Isles and outside of summer (Figure 5). Metadata accuracy (43.7%) was most related to temperature. This is demonstrated by a 10% percentage point decrease in accuracy when temperature was removed. Where both temperature and day-of-year data was available, the temperature data appears to be used more (10% and 0.2% decreases respectively). It is not possible to determine whether this is because temperature is simply more relevant to ladybirds than date, or whether this is an artefact of the different lengths of the metadata vectors. When day-of-year was randomised in the primary metadata model, top-3 accuracy declines by 4.5% points. Within temperature, the model appeared to be making more use of the weekly temperature data (2-10 weeks before the record), where randomisation caused an 8.1% decrease than the more proximate daily records for the preceding fortnight (−5.4%). The remaining metadata components had smaller influences on overall top-3 accuracy: randomising habitat data led to a 2.8% decrease while randomising recorder experience led to a 2.1% decrease.

**Figure 5.**
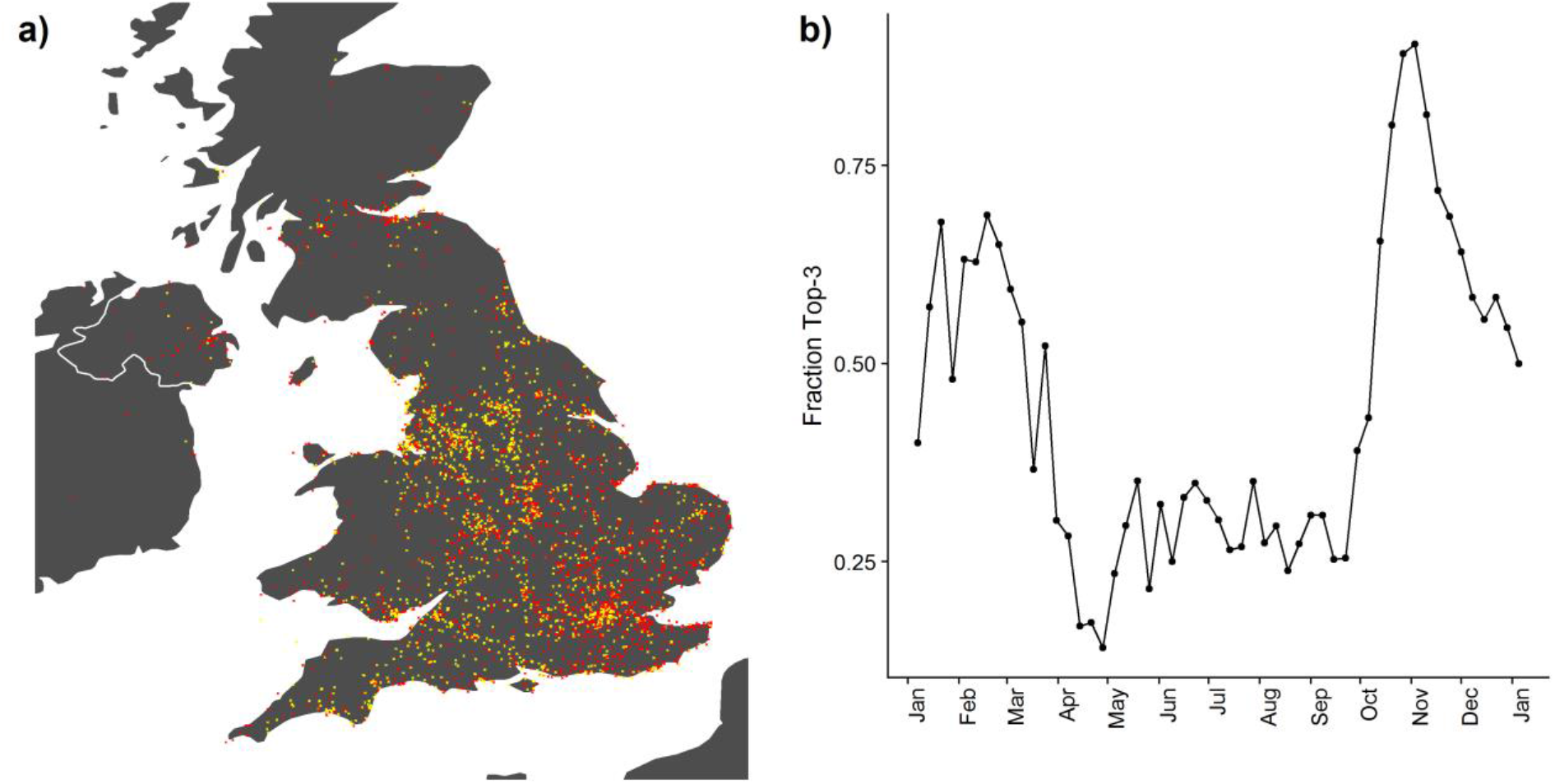
Distribution of records accurately (top-3) predicted solely from a derived metadata model. a) Spatial distribution of accuracy, showing decreased accuracy in the south-east. Accurate predictions are shown in yellow, incorrect in red, b) Weekly fraction of accurate metadata identifications through the year showing strong seasonal variation in accuracy with a particular peak in mid-autumn.

These overall results are highly influenced by the dominant species (particularly the Harlequin ladybird) in the test set, masking variation in decline in accuracy on a per-species level (SI Table S2). The apparent importance of each metadata component appears to align with ecological expectations. The five species with greatest decline in accuracy when habitat is randomised are all considered habitat specialists (Roy & Brown, 2018): *Coccinella undecimpunctata* (dunes), *Anatis ocellata* (conifers), *Tytthaspis sedecimpunctata* (grassland and dunes), *Subcoccinella vigintiquattuorpunctata* (grassland), and *Aphidecta obliterata* (conifers). Similarly, the randomisation of location had the greatest effect on the localised species (Figure S1). The top three most affected were: *Aphidecta obliterata (*frequently reported in Scotland), *Scymnus interruptus* (South-East England) and *Coccinella undecimpunctata* (coastal). By contrast, the Seven-Spot ladybird, a widespread and generalist speices was poorly identified by the metadata model and showed a minimal response to randomisation. The species affected most by the randomisation of temperature was *Propylea quattuordecimpunctata*, with the common name of the ‘dormouse’ ladybird (Roy and Brown 2018, p.112) because of its known late emergence.

The randomisation of recorder experience had the greatest impact on *Scymnus interruptus*. This was the only ‘inconspicuous’ ladybird in our dataset, which inexperienced recorders may not even realise is a ladybird (see Figure 2g). There was also a 10% decrease in the identification of Harlequin ladybirds when recorder experience was randomised. Novice recorders are notably more likely to record Harlequin ladybirds than more experienced recorders. The first record submitted by a new recorder is a Harlequin ladybird 57.4% of the time, which rapidly declines to 38% by the 10^th^.

## Discussion

The use of metadata within computer vision models considerably improves their reliability for species identification. This exciting finding has implications for biological recording, demonstrating the potential to use innovative approaches to assist in processing large occurrence datasets accrued through mass participation citizen science. Basic primary metadata is straightforward to incorporate within machine learning models and, since this information is already collected alongside the biological records, can be widely adopted.

### Interpretation of results

The notable gain in accuracy of the combined multi-input model compared to the ensemble model is consistent with the model learning to interpret the image based on the metadata. This is evidence that metadata can provide further gains beyond simply filtering the potential species list (Wittich et al., 2018). While it is not possible to determine exactly what interpretations the artificial intelligence is making, we can discern plausible scenarios. In autumn, ladybirds select suitable overwintering sites and enter dormancy through the adverse months (Roy & Brown, 2018). Each species exhibits a specific preference in overwintering habitat. Harlequin ladybirds favour buildings, leading to a high proportion of submitted records from inside homes of Harlequin ladybirds in the autumn as they move inside to overwinter (Roy et al., 2016). Submitted images of ladybirds exhibiting this behaviour are often poor-quality showing ladybirds at a distance nestled in crevices (Figure 2). The high accuracy of the metadata model during autumn suggests it has learnt (as expert human verifiers have) that a poor-quality image with a pale background during the autumn is very likely a Harlequin ladybird.

Our results likely represent a lower bound on the potential improvements that can be leveraged from metadata for identifying challenging species. Although British ladybirds have distinct ranges, activity periods and habitat (Comont et al., 2012; Roy & Brown, 2018) many are relatively cosmopolitan and can be observed as adults for large parts of the year. Classification models where focal species are more localised in time, space or habitat, or alternatively if the domain of the model is larger (for example North America, Berg *et al*. 2014), may expect to see larger gains through including metadata.

Determining how deep learning models make decisions is complex (Goodfellow et al., 2016). Multiple interwoven contributing factors combine to produce a result, much akin to human decisions. The nature of metadata means much of the gain likely comes from ruling species out rather than positively identifying them, which makes the interpretation of ‘accuracy’ metrics even more challenging. Our randomisation analysis to determine the features used by the metadata model can only be a rough guide to the basis of decisions. The randomisation process will represent the pre-existing imbalance of our dataset and will produce illogical combinations of metadata, such as hot temperatures during the winter, or coastal habitat within inland areas. Nonetheless, it does show evidence that the model operates along similar lines to expert identifiers. Where certain aspects of information are lost, this translated into inaccuracies in species for which that information is relevant. This is aligned with the results of Miao et al. (2018) who found that their image recognition tool for savanna mammals also used similar features to humans to identify species. Equally, for widespread and generalist species, metadata is not able to contribute to the accuracy. For instance, the identification of Seven-spot ladybird is essentially unchanged by the inclusion of metadata.

In theory, given enough records, a deep-learning model would be able to infer the information content of the cross-referenced database based only on primary metadata. For example, a neural network could learn to identify a set of location coordinates with a high likelihood of a given species, without knowing that those coordinates contained favoured habitat, simply because the species is frequently recorded at these locations in the training dataset. In this respect, the inclusion of derived metadata could be considered a feature extractor technique that interprets the primary metadata, rather than providing additional information. In practice, the level of data required to internally reconstruct sufficient mapping purely from primary metadata would be very high, particularly when the features are very high resolution (Tang et al., 2015). A core challenge for automated species identification is the long tail of species for which there are very sparse records (Van Horn et al., 2017), for which the advantage of including derived metadata is likely to be considerably larger than for frequently recorded species.

### Further Improvements to Model

The design and training of deep learning models is an art rather than an exact science (Chollet & Allaire, 2018). There are likely to be opportunities for improvement in overall accuracy for each of our models. Our image-only accuracy levels (48.2%) were below that attained on other ecological datasets, though citizen scientists’ images of ladybirds have been previously identified as posing a particular challenge for computer vision systems (Van Horn et al., 2017). For example, 67% accuracy was established as a baseline on the diverse iNaturalist image competition dataset (Van Horn et al., 2017), while competition winners were able to reach 74%.

Practically, incorporating metadata into neural networks need not introduce considerably more effort. Metadata is substantially simpler to process than image data and did not appear to add significantly to the training time. Compared to the very deep convolutional networks needed to interpret images, metadata can be processed with a small number of densely connected layers. Our tests with much larger or deeper networks did not lead to further gains. The number of parameters in our metadata models were several orders of magnitude smaller than the image model and could be trained in a matter of seconds per epoch. However, there are small additional design overheads in constructing a multi-input neural network compared to an image-only approach. There now exist user-friendly ‘automatic learning’ software that can generate a computer vision model given only a set of labelled images. In contrast, currently available support for multi-input models is comparatively lacking and requires direct specification of the model architecture as well as data manipulation pipelines to combine disparate information sources. Fortunately, tools such as the *keras* R package (Allaire & Chollet, 2019) provide straightforward frameworks for multi-input models that are well within the reach of ecologists without a formal computational science background. We have also shared our code (SI) to help others make use of this methodology.

We have demonstrated the improvement gained through the use of metadata. Further improvements could likely be made through instigating test-time augmentation where multiple crops or rotations of an image are presented to the classifier, ensembling multiple models, and increasing the size of the dataset through supplementary images and historical records (Chollet & Allaire, 2018). Our approach to augmenting metadata (adding Gaussian noise to each element) was relatively basic and more targeted approaches to generating additional synthetic training data (Chawla et al. 2002) could lead to better results.

The overall accuracy of a species classifier can be considerably enhanced by incorporating a prior likelihood of each species’ relative frequency. Approaches that allow the model to directly learn the relative frequencies of the species could attain even higher overall accuracy. However, in contrast to improvements discussed in the previous paragraph this would significantly reduce the accuracy for rarely observed species. A model that only learnt to accurately distinguish between Harlequin and Seven-spot ladybirds (that constitute the majority of records) could attain an accuracy of 70%, but this would be of limited applied use.

The challenge of species identification has in the past attracted computer scientists who can view species identification as an interesting example of large real-world labelled datasets (Weinstein 2018). Open competitions such as the annual iNaturalist (Van Horn et al., 2017) and LifeCLEF competitions (Goëau, Bonnet, & Joly, 2017) have spurred considerable improvements in identification accuracy. Including metadata in these datasets (such as the PlantCLEF 2019 competition) could lead to considerable improvements. However, any release of metadata must consider the geoprivacy of citizen scientists and potential risk to endangered species. Due consideration of the appropriate resolution of location data, and the identifiability of individuals in any data publicly released is essential.

### Transferability of models including metadata

The inclusion of metadata in an automatic identification tool will influence its transferability to new contexts. With all machine learning approaches, any automatic identification process is only as good as the extent and scope of the training data used. A model that has been trained on the location of UK records would need to be retrained for use in continental Europe, whereas an image-only model could be expected to be at least somewhat useful in both contexts. As such, a model trained on derived metadata such as habitat types or local weather may be more transferable than one trained on coordinates and specific dates. A focussed appreciation of the domain a model will be applied to is essential. Transferability will be critical for expanding from well-studied areas (such as UK), to understudied areas where there is great potential for citizen science to fill gaps in knowledge (Pocock et al., 2018).

Transferability of models can be a challenge even within a region since records generated through unstructured broad-based citizen science are distinctive from those generated by committed amateur recorders, structured citizen science projects or professional surveys (Boakes et al., 2016). Submitted records are the result of interactions between human behaviour and species ecology (Boakes et al., 2016). Highly visited sites may show an over-abundance of common species that are new to citizen scientists with relatively limited experience. In our dataset, uploaded records of ladybirds correlate strongly with the first appearance of species and news reports of invasive species (T. A. August *unpublished data*).

Our choice of what contextual data to include was guided by our knowledge of variables that are likely to influence ladybirds in the British Isles. For more taxonomically diverse tools, it would be beneficial to use a wider a range of derived metadata variables. This could include more diverse weather information, climate maps, and topography. We did not include species range maps (Roy, Brown, Frost, & Poland, 2011) in this study since most (>90%) records came from areas within the range of 15 out of the 18 focal species considered in this study. Binary species range maps cannot account for the relative frequency of species across a region, but this can be learnt by a deep learning network provided with location data of records. Although range maps could be informative within models with a wide spatial scope or for highly localised species, they are comparatively verbose to encode for in deep learning networks. When using a model to identify large numbers of species, the intersection or otherwise of a record with each species range map may need to be encoded in a separate variable.

This greatly increases the length of the metadata vector associated with each record and it could become challenging for models to identify relevant information. Although deep learning networks have the potential to effectively ignore data that is not relevant, there is the potential to slow the fitting procedure if too much irrelevant information is presented. Where accurate species range map data is available (and may impart additional information beyond that contained in the training set of records), an approach that combines machine learning with a range-map based shortlist may be the most useful (Wittich et al., 2018).

### Conclusions

Identification of insects poses a considerable challenge for computer vision (Martineau et al., 2017). Insect diversity is extraordinarily large – as an example, there are over 6000 ladybird species worldwide (Roy and Brown 2018), most of which do not have accessible labelled images. For difficult challenges, such as species identification in the field, the optimal solutions will involve humans and artificial intelligence working in tandem (Trouille, Lintott, & Fortson, 2019). Our results demonstrate the potential for considerable improvement in the accuracy of automatic identification when incorporating contextualisation information directly within the model. This is also likely to apply to passive acoustic monitoring tools (Gibb, Browning, Glover-Kapfer, & Jones, 2019) too. Researchers building automatic identification methods will benefit from training models to place images in context, just as a human naturalist would, to best unlock the potential of artificial intelligence in ecology.

## Acknowledgments

Our thanks to Mark Logie for assistance accessing the iRecord database, Colin Harrower for species range maps, and to the UK Ladybird Survey volunteer recorders who generated the dataset the work is based upon. This work was supported by the Natural Environment Research Council award number NE/R016429/1 as part of the UK-SCAPE programme delivering National Capability. JCDT was funded for this project through a NERC Innovation placement linked to the Oxford Environmental Doctoral Training Program (NE/L002612/1).

## Authorship Statement

JCDT built the models and analysed the results, based on an initial idea and design of TAA and through discussions with HER and TAA. JCDT wrote the first manuscript draft and all authors contributed critically to revisions.

## Data Accessibility

R code used to build models and analyse results is available at https://github.com/jcdterry/LadybirdID_Public and summarised in SI 2. Records were accessed from the UK Biological Records Centre (www.brc.ac.uk).

