## Supplementary Information for "Thinking like a naturalist: enhancing computer vision of citizen science images by harnessing contextual data"

Supplementary Figures, Tables and Appendices for *Thinking like a naturalist: enhancing computer vision of citizen science images by harnessing contextual data*

J. Christopher D. Terry, Helen E. Roy & Tom A. August

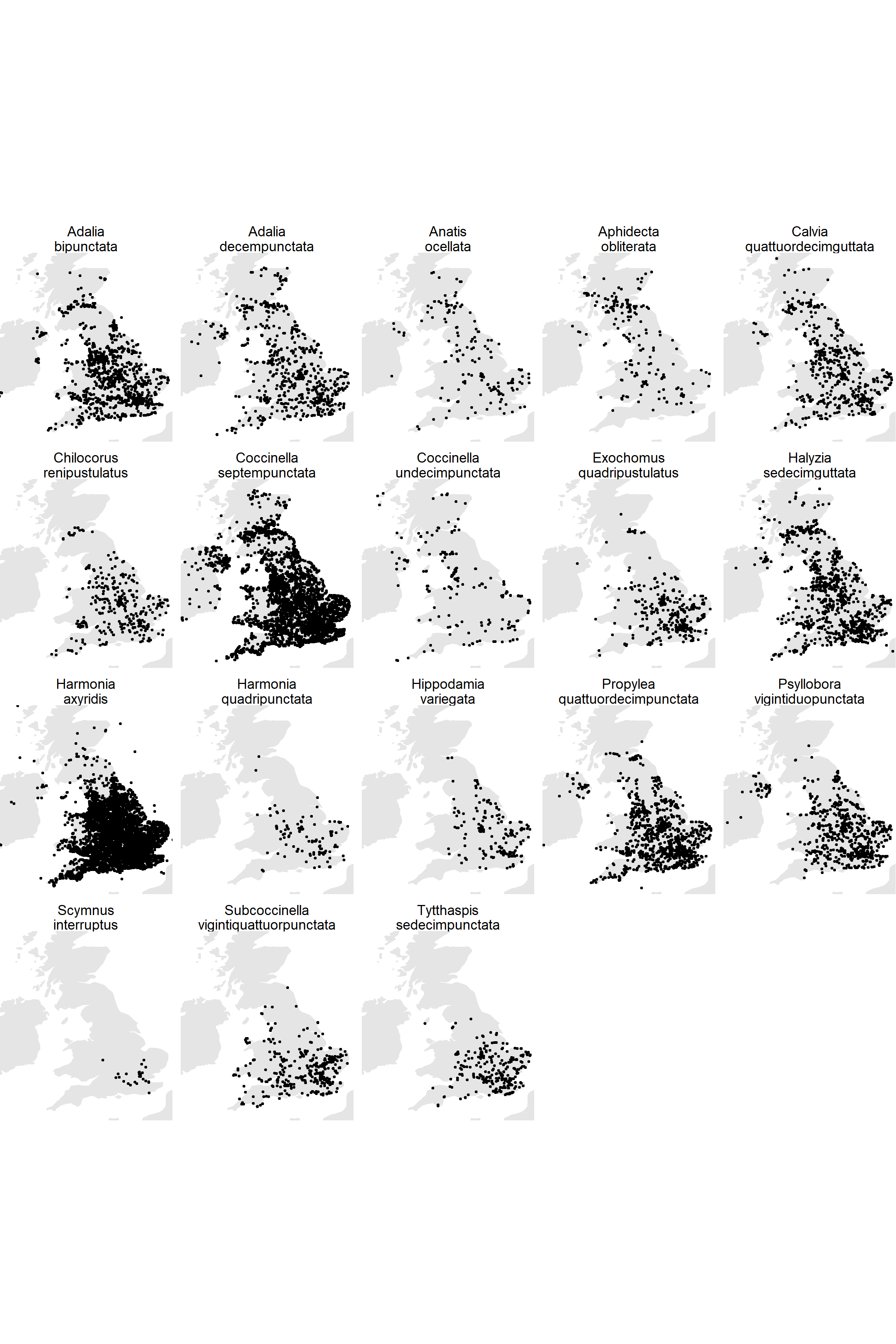

**Figure S1.** Spatial distribution of records for each of the 18 ladybird species that the tool seeks to identify. Most species have been reported across large areas of the country, while a few such as *Scymnus interruptus* and *Tytthaspis sedecimpunctata* are more localised.

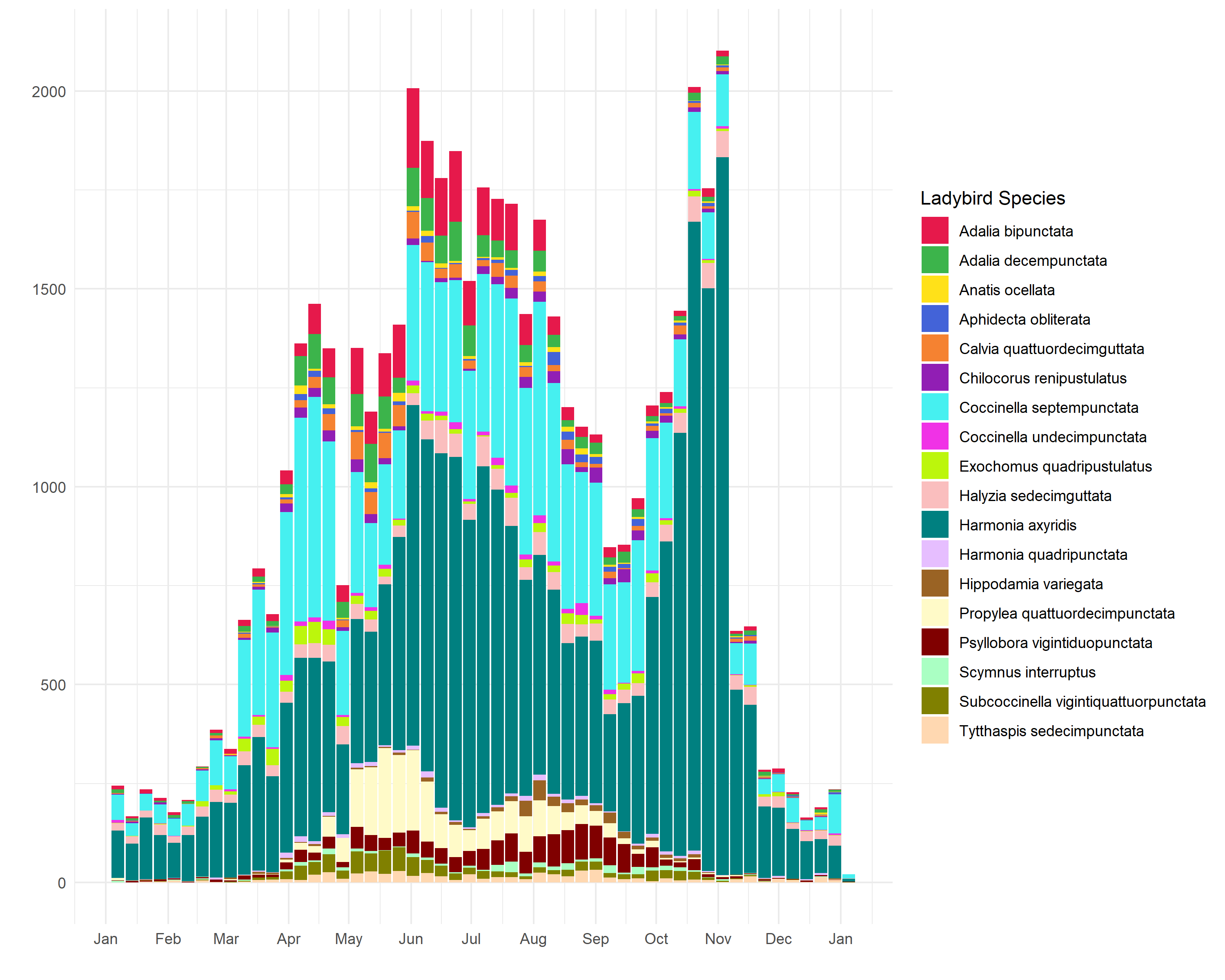

**Figure S2.** Distribution of ladybird records grouped by week throughout the year. The distribution is approximately tri-modal with peaks in spring when the first ladybirds emerge, in summer as abundance reaches its peak and October when certain species (especially Harlequin ladybirds) enter houses in order to overwinter.

**Table S2** Percentage decline in accuracy as categories of metadata are randomised

| **Name** | **Baseline**  **Accuracy** | **Day-of-Year** | **Habitats** | **Location**  **(Latitude and longitude)** | **Recorder**  **Experience** | **All Temperatures** | **Weekly Temperatures** | **Daily Temperatures** |
| --- | --- | --- | --- | --- | --- | --- | --- | --- |
| Adalia  bipunctata | 47.6 | -0.1 | 10.7 | 5.4 | 1.9 | 41.2 | 36.5 | 17.5 |
| Adalia  decempunctata | 21.5 | 4.8 | 5.9 | 42.0 | 5.9 | 37.2 | 29.8 | 4.3 |
| Anatis  ocellata | 41.3 | -1.6 | 22.6 | 45.2 | 6.5 | 38.7 | 25.8 | 22.6 |
| Aphidecta  obliterata | 67.4 | 0.0 | 12.5 | 64.8 | 0.0 | 7.8 | 7.0 | 4.7 |
| Calvia  quattuordecimguttata | 24.9 | -0.8 | -1.5 | 15.9 | 5.3 | 35.6 | 15.9 | 10.6 |
| Chilocorus  renipustulatus | 34.4 | -4.1 | 9.8 | 9.0 | -6.6 | 23.0 | 16.4 | 19.7 |
| Coccinella  septempunctata | 14.8 | 3.4 | 0.1 | 0.3 | 3.3 | 32.6 | 44.2 | 41.1 |
| Coccinella  undecimpunctata | 44.1 | -1.2 | 27.9 | 59.3 | 5.8 | 4.7 | 12.8 | 16.3 |
| Exochomus  quadripustulatus | 48.0 | 0.0 | 5.3 | 29.2 | 8.6 | 26.3 | 24.9 | 14.8 |
| Halyzia  sedecimguttata | 39.6 | -0.7 | 8.9 | 15.8 | 0.9 | 33.6 | 11.6 | 18.5 |
| Harmonia  axyridis | 61.1 | 0.6 | 7.0 | 8.9 | 9.0 | 15.6 | 11.4 | 5.9 |
| Harmonia  quadripunctata | 34.6 | 0.0 | 2.2 | 35.6 | 11.1 | -4.4 | 28.9 | 17.8 |
| Hippodamia  variegata | 53.1 | -1.1 | 3.2 | 36.6 | 6.5 | 36.6 | 32.3 | 7.5 |
| Propylea  quattuordecimpunctata | 50.7 | 0.4 | -0.7 | 5.6 | 1.8 | 49.4 | 34.9 | 22.8 |
| Psyllobora  vigintiduopunctata | 41.7 | 0.5 | -0.3 | 13.7 | -1.9 | 37.2 | 33.2 | 32.3 |
| Scymnus  interruptus | 85.6 | 0.0 | 2.8 | 61.7 | 39.3 | 10.3 | 26.2 | 16.8 |
| Subcoccinella  vigintiquattuorpunctata | 53.5 | -2.0 | 20.1 | 33.3 | 12.0 | 43.0 | 12.0 | 10.0 |
| Tytthaspis  sedecimpunctata | 53.3 | 0.6 | 20.8 | 48.2 | 11.3 | 23.8 | 11.3 | 16.7 |

Appendix 1: Metadata scaling factors.

Machine learning models tend to converge faster when given small input values. For this reason we scaled the majority of our inputs:

- RGB images values were normalised from [0:255] to the range [-1:1] by ((x/255)-0.5)*2
- Day of year was scaled from 1:366 range to [0:1] by dividing by 366
- Recorder experience ranged from 1 to 1047. This was scaled by taking the base-10 logarithm.
- Habitat proportion already ranged between 0:1. It was left unrescaled since there are batch normalisation layers after the input to the metadata model.
- Longitude, Latitude, weekly max temperature averages and daily max temperature measurements were each normalised by subtracting the corresponding mean and dividing by corresponding SD:

| **Metadata** | **Mean** | **SD** |
| --- | --- | --- |
| Temperature daily | 16.25722 | 5.128868 |
| Temperature Weekly | 15.32672 | 5.18819 |
| Latitude | 52.53597 | 1.298351 |
| Longitude | -1.433536 | 1.569204 |

Appendix 2: Model Structures

‘keras’ R code to build the models is available on github https://github.com/jcdterry/LadybirdID_Public. However, since that is complicated by additional code for data manipulation and sequential model fitting, we detail the core model structures here:

*Image Only Model:*

```
conv_base_IRv2 <- application_inception_resnet_v2(weights = 'imagenet',
                                                  include_top = FALSE,
                                                  input_shape = c(299,299,3),
                                                  pooling ='max')

Image_Predictions<- conv_base_IRv2$output%>%
  layer_dropout(rate=0.3,
                name = 'Image_Base_Dropout_1') %>%
  layer_batch_normalization(name = 'image_normalisation')%>%
  layer_dense(units = 256,
              activation = 'relu',
              kernel_regularizer =  regularizer_l2(l = 0.0001),
              name = 'BaseTop1')   %>%
  layer_dropout(rate=0.3,
                name = 'Image_Base_Dropout_2') %>%
  layer_dense(units =18,
              activation = 'softmax',
              name = 'main_output')

Image_Model <- keras_model(inputs = conv_base_IRv2$input,
                           outputs = Image_Predictions)

Image_Model %>% compile(  loss = "categorical_crossentropy",
                          optimizer = optimizer_adam(lr = lr$IM1),
                          metrics='categorical_accuracy')
```

*Metadata only model*

Primary metadata only:

```
meta_input = layer_input(shape=  3, name='metadata_input')

meta_predictions<- meta_input %>%
  layer_gaussian_noise(stddev = 0.1,
                       name = 'meta_noise')%>%
  layer_batch_normalization(name= 'meta_input_normalisation')%>%
  layer_dense( units=16,
               activation = "relu",
               name = 'meta1',
               kernel_regularizer = regularizer_l2(0.00001) ) %>%
  layer_dropout(0.3, name = 'meta_dropout2')%>%
  layer_dense( units = 16,
               kernel_regularizer =  regularizer_l2(0.00001),
               activation = 'relu',
               name = 'meta2' )%>%
  layer_dropout(0.3, name = 'meta_dropout3')%>%
  layer_dense(units = 18,
              activation = 'softmax', name = 'meta_output')

Meta_Model<- keras_model(inputs = meta_input, outputs = meta_predictions)

Meta_Model %>% compile(  loss = "categorical_crossentropy",
                         optimizer = optimizer_adam(lr = lr$M1),
                         metrics = "categorical_accuracy")
```

Primary and Secondary metadata:

```
meta_input = layer_input(shape=  47, name='metadata_input')

meta_predictions<- meta_input %>%
  layer_gaussian_noise(stddev = 0.2,
                       name = 'meta_noise')%>%
  layer_batch_normalization(name= 'meta_input_normalisation')%>%
  layer_dense( units=64,
               activation = "relu",
               name = 'meta1',
               kernel_regularizer = regularizer_l2(0.0001) ) %>%
  layer_dropout(0.2, name = 'meta_dropout2')%>%
  layer_dense( units = 64,
               kernel_regularizer =  regularizer_l2(0.0001),
               activation = 'relu',
               name = 'meta2' )%>%
  layer_dropout(0.2, name = 'meta_dropout3')%>%
  layer_dense(units = 18,
              activation = 'softmax', name = 'meta_output')

Meta_Model<- keras_model(inputs = meta_input, outputs = meta_predictions)

Meta_Model %>% compile(  loss = "categorical_crossentropy",
                         optimizer = optimizer_adam(lr = lr$M2),
                         metrics = "categorical_accuracy")
```

*Combined Model With Image and Metadata*

```
base_image_model<-load_model_hdf5('Image_Model_S2')
base_meta_model<- load_model_hdf5('Meta_Model_Secn'))

base_image_model_decap<-(base_image_model %>% get_layer('BaseTop1'))$output
base_meta_model_decap<-(base_meta_model %>% get_layer('meta2'))$output

BothTogetherOutput<-list(layer_batch_normalization(base_image_model_decap,
                                                   name = 'Image_Normaliser'),
                         layer_batch_normalization(base_meta_model_decap,
                                                   name = 'Meta_Normaliser'))%>%
  layer_concatenate( name='LinkMeta_BaseTop')%>%
  layer_dropout(rate=0.4,
                name= 'Post-ConCat_Dropout')%>%
  layer_dense(units = 64,
              activation = 'relu',
              kernel_regularizer =  regularizer_l2(l = 0.001),
              name = 'Combo1')%>%
  layer_dropout(rate=0.4,
                name= 'Final_PostConCat_Dropout')%>%
  layer_dense(units = 18,
              activation = 'softmax', name = 'Combined_output')

MultiInputModel_1<-keras_model(inputs = c(base_image_model$input,
                                          base_meta_model$input),
                               outputs = BothTogetherOutput)

MultiInputModel_1 %>% compile(  loss = "categorical_crossentropy",
                                optimizer =  optimizer_adam(lr = lr$B1),
                                metrics='categorical_accuracy')
```
